# The TPLATE complex mediates membrane bending during plant clathrin-mediated endocytosis

**DOI:** 10.1101/2021.04.26.441441

**Authors:** Alexander Johnson, Dana A Dahhan, Nataliia Gnyliukh, Walter A Kaufmann, Vanessa Zheden, Tommaso Costanzo, Pierre Mahou, Mónika Hrtyan, Jie Wang, Juan Aguilera-Servin, Daniël van Damme, Emmanuel Beaurepaire, Martin Loose, Sebastian Y Bednarek, Jiri Friml

**Affiliations:** Institute of Science and Technology (IST Austria), 3400 Klosterneuburg, Austria; UW-Madison, Biochemistry, 215C HF DeLuca Laboratories, Madison, WI, 53706; Laboratory for Optics and Biosciences, Ecole Polytechnique, CNRS, Inserm, IP Paris, Palaiseau, France; Ghent University, Department of Plant Biotechnology and Bioinformatics, Technologiepark 71, 9052 Ghent, Belgium; VIB Center for Plant Systems Biology, Technologiepark 71, 9052 Ghent, Belgium

## Abstract

Clathrin-mediated endocytosis in plants is an essential process but the underlying mechanisms are poorly understood, not least because of the extreme intracellular turgor pressure acting against the formation of endocytic vesicles. In contrast to other models, plant endocytosis is independent of actin, indicating a mechanistically distinct solution. Here, by using biochemical and advanced microscopy approaches, we show that the plant-specific TPLATE complex acts outside of endocytic vesicles as a mediator of membrane bending. Cells with disrupted TPLATE fail to generate spherical vesicles, and *in vitro* biophysical assays identified protein domains with membrane bending capability. These results redefine the role of the TPLATE complex as a key component of the evolutionarily distinct mechanism mediating membrane bending against high turgor pressure to drive endocytosis in plant cells.

**One Sentence Summary:** While plant CME is actin independent, we identify that the evolutionarily ancient octameric TPLATE complex mediates membrane bending against high turgor pressure in plant clathrin-mediated endocytosis.

## Main Text

Clathrin-mediated endocytosis (CME) is a critical eukaryotic cellular process that regulates a wide range of physiological processes, for example, mediating the internalization of receptors and transporters (*1*). During CME, a small area of the plasma membrane (PM) invaginates into the cell forming a spherical vesicle against intracellular forces that oppose membrane deformation, like turgor pressure. The mechanisms driving this process in mammalian and yeast systems have been the subject of extensive study for the better part of 5 decades, which has led to the identification of key proteins that provide the force required to overcome these forces (*2, 3*). Critically, actin has been established to be essential for membrane bending in systems with high turgor pressures (*2, 4*). In stark contrast, plant CME characterization is in its infancy. Indeed, it had been postulated for many years that due to the extreme levels of turgor pressure in plants, CME was physically impossible in most plant cells (*5*). However, while it is now well established that CME does occur *in planta*, and plays key developmental and physiological roles (*6*), the machinery and mechanisms that drive CME against the unique biophysical properties of plant cells are yet to be clearly identified. For example, it had long been thought that plant CME relies on actin to overcome the extreme turgor pressure, however it has recently been demonstrated that plant CME is independent of actin, highlighting that plants have evolved a distinct solution to bending membranes against high turgor pressures (*7*). A further mechanistic divergence of plant CME is manifested by the presence of the octameric TPLATE complex (TPC), where all 8 members share the same localizations and dynamics at sites of plant CME, and is essential for both CME and plant survival (*8, 9*). While this complex is conserved in some biological systems, for example *Dictyostelium*, it is notably absent from mammalian and yeast genomes (*8, 10, 11*). Based on static interaction and localization data, the TPC has been proposed to be a classical endocytosis adaptor protein; chiefly acting to bind cargo in the clathrin-coated vesicle (CCV), driving the coat assembly, and has thus been predicted to localize beneath the clathrin coat (*8, 12*).

However, recent total internal reflection fluorescence microscopy (TIRF-M) of CME events *in planta* suggested that once the endocytic CCV departs from the PM, TPLATE dissociates from the CCV prior to the loss of clathrin (*7*). This did not support the proposed adaptor functions of TPLATE, which, as assumed to be localized under the clathrin coat, should have an equivalent/slower dissociation relative to clathrin. Furthermore, the classical adaptor, AP2, and TPLATE have different dynamics at co-localized foci on the PM (*8*). To further assess and quantify the dissociation of TPLATE from the CCV, we analyzed CME events marked with Clathrin light-chain 2 (CLC2-tagRFP) and TPLATE-GFP obtained using a spinning disk microscope equipped with a sample cooling stage (Fig. 1A) (*9*). As this imaging modality provides an increase in the illumination volume of the cytoplasm compared to TIRF-M, and cooling the sample slowed the dynamics of cellular processes (*9*), it allowed a more precise visualization of the dissociation sequence of proteins from the CCV once it is freed from the PM (Fig. S1A). This resulted in kymographs of individual CME events with a visible lateral divergence of fluorescence signals at the end of the CME events (Fig. 1B), which represent the movement of CCVs once departed from the PM. These ‘departure traces’ were analyzed to establish the sequence of TPLATE and CLC2 dissociation. We found that the significant majority of visible departure traces displayed differential dissociation of these proteins; critically where TPLATE departed the CCV before CLC2 (Fig 1C., Fig S1B.). This argued against the model that TPC is entrapped within the clathrin coat of the CCV. To further query the nature of the association of the TPC with CCVs, we performed western blot analysis of purified CCV preparations from plant cells. While we found an enrichment of the clathrin isoforms and the canonical adaptor AP2 in the purified CCV fraction, TPLATE was not enriched (Fig 1C-D). The depletion of TPLATE was observed in parallel with a similar lack of enrichment of dynamin related protein 1c (DRP1c), which functions outside the CCV (*13*). Together, this demonstrates that TPLATE is more loosely associated with CCVs than clathrin coat proteins and not incorporated within CCV structures as previously predicted.

**Fig. 1.**
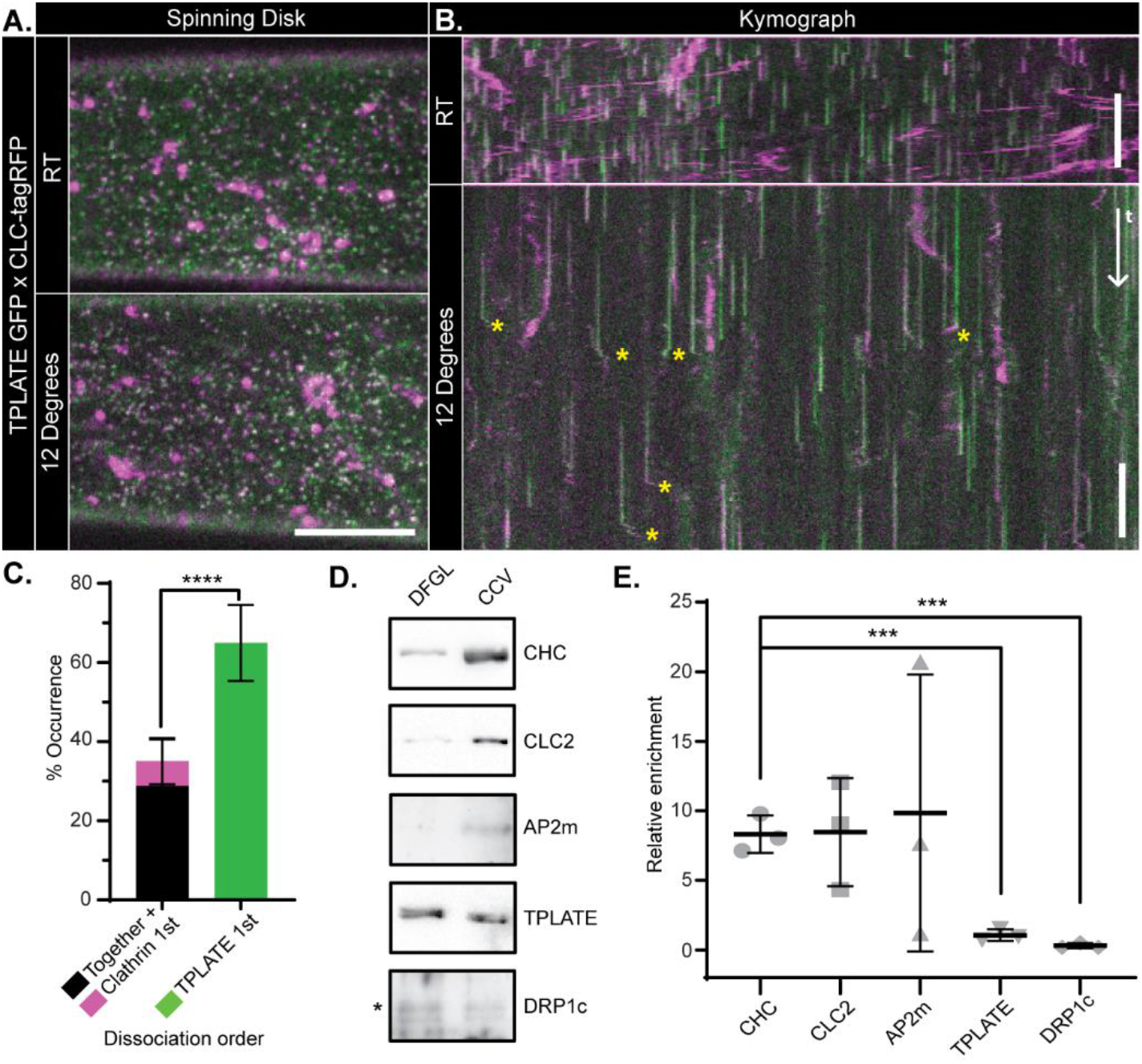
TPLATE is only loosely associated with CCVs. A) Example spinning disk images and B) kymographs of *Arabidopsis* hypocotyl epidermal cells expressing TPLATE-GFP (green) and CLC2-TagRFP (magenta) at either room-temperature or 12°C. Yellow asterisks note example departure traces where CCVs are visible after dissociation from the PM. C) Quantification of departure traces based on the order of departure (Fig. S1B). N = 14 cells from independent plants, 258 departure traces. ****p < 0.001, T-Test. D) Representative western blots of endocytosis proteins during CCV purification and E) quantification of proteins in the CCV fraction relative to an earlier purification step (DFGL). N = 3 independent CCV purifications. *** p > 0.01, T-tests compared to Clathrin heavy chain (CHC). Plots, mean ± SD. Scale bars, A, 5 µm; B, 60 s.

To precisely determine the localization of TPLATE at CME events we used structured illumination microscopy (SIM) to examine live plants expressing TPLATE-GFP and CLC2-TagRFP. We observed that TPLATE, in addition to presenting as individual foci, often appeared as crescent shaped or ring structures (Fig. 2A and Fig. S2A). This was in contrast to CLC2, which was always found as discrete foci on the PM or large structures representing trans-Golgi/ early endosomes (*7*). We found that ∼68% of TPLATE co-localized with CLC2 foci, agreeing with previously published results (*8*). Upon closer inspection of the super resolved co-localized events, we found that ∼17% of co-localization events showed that TPLATE formed a ring or crescent around the CLC2 foci whereas the inverse arrangement was never observed, suggesting that TPLATE localizes outside of the clathrin coat assembly during CME. In further support of this, when we examined plants expressing TPLATE-GFP and AP2A1-tagRFP we found a similar pattern, where ∼18% of colocalized events showed TPLATE surrounding AP2 in ring and crescent patterns (Fig. S2A). To gain further insight into the TPLATE ring arrangements, we visualized the dynamics of TPLATE on the PM using TIRF-SIM. We observed that TPLATE first appeared as a spot, and over time form a ring, which then closed back to a spot before disappearing (Fig. 2C and Movie 1). This suggests that at the beginning of the CME event, TPLATE and clathrin are both present in an area below the resolution of SIM, but as the CCV grows in diameter (above the resolution limit), TPLATE is excluded from the invaginating CCV coat formation and is localized at the rim around the CCV. This confirms that TPLATE is localized at the periphery of CCV events, rather than within the invaginating clathrin coat.

**Fig. 2.**
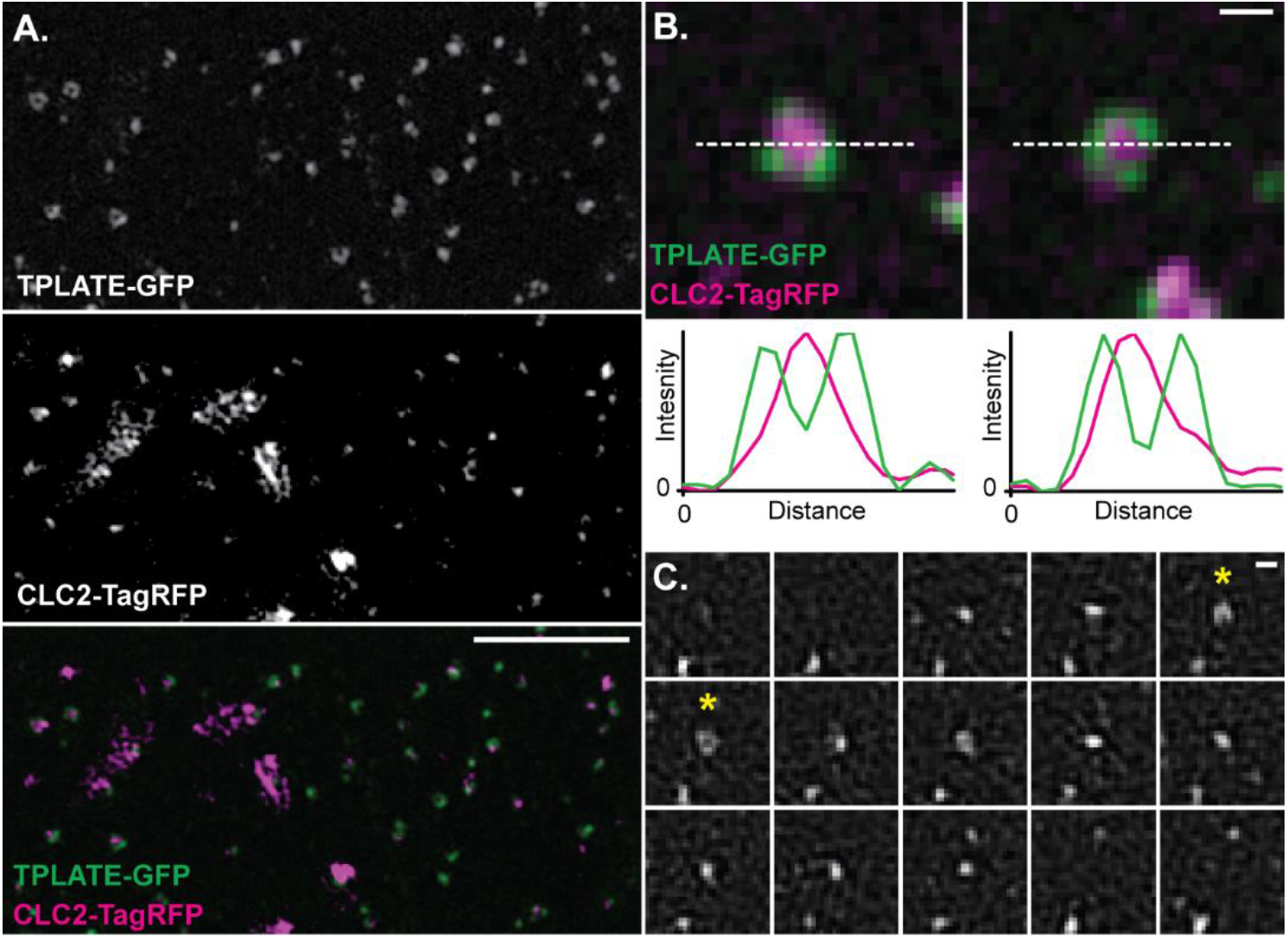
The TPLATE complex is localized at the rim of CME events. A) Representative 3D-SIM image of an *Arabidopsis* root epidermal cell expressing TPLATE-GFP and CLC2-TagRFP. B) Examples of individual endocytosis structures and line plots (white dotted line) of their fluorescent intensities. C) TIRF-SIM example of TPLATE-GFP dynamics in an Arabidopsis root epidermal (See Movie 1 for a larger field of view). Asterisks note when a ring structure is formed. Frame interval is 5 seconds. Scale bars, A, 3 µm; B and C, 200 nm.

In mammalian and yeast CME, the endocytic proteins which are localized at the rim of the CME events are implicated in membrane bending, for example Eps15/Ede1, Epsin and Fcho/Syp1 (*14-17*). Therefore, based on the spatial and temporal profile of TPLATE, and the ability of distinct domains of TPC to bind directly to the PM (*18, 19*), we hypothesize that components of the TPC are critical for membrane bending during plant CME. To test this notion, we looked directly at CCVs in plant cells subjected to TPC disruption. For these studies, we used the inducible TPLATE loss-of-function mutant WDXM2; where *tplate* mutant plants are complemented with a genetically destabilized version of TPLATE that after heat shock results in the total aggregation of TPLATE away from the PM and blocks CME (*20*). First, we further confirmed that the heat shock treatment had no significant effect on the efficiency and kinetics of CME in plants by examining FM4-64 uptake and the dynamics of single CME events (Fig. S3). To achieve the spatial resolution required to look at the shape of individual CCVs, we used scanning electron microscopy (SEM) to examine metal replicas of unroofed protoplasts made directly from wild type or WDXM2 roots. Under control conditions in WDXM2 cells, we observed that the majority of clathrin structures were spherical (i.e., fully invaginated CCVs). In contrast, in cells in which the TPC was disrupted, we observed many flat clathrin structures (Fig. 3A), which were never observed in control conditions. To quantify the effect of TPC disruption, we determined the shape of the clathrin structures by measuring the area and average intensity (a proxy for CCV curvature (Fig. S4A) (*21*)) of each clathrin structure visualized and classified these shapes into 4 categories; ‘small and round’ (the fully invaginated CCVs), ‘small and flat’ (where curvature generation had failed), ‘large and round’, and ‘large and flat’ (clathrin plaques) (Fig. 3C). We found that the heat shock had no effect upon CCV formation in wild type cells (Fig. S4B and Fig. 3D), where the majority of clathrin structures were found as ‘small and round’ (87-96%). This population in WDXM2 cells under control conditions was 86%, but following TPC disruption, this decreased to 24% and the ‘small and flat’ population increased to 58% (compared to <7% in all other tested conditions). These results indicated that the TPC is required to generate curved clathrin structures.

**Fig. 3.**
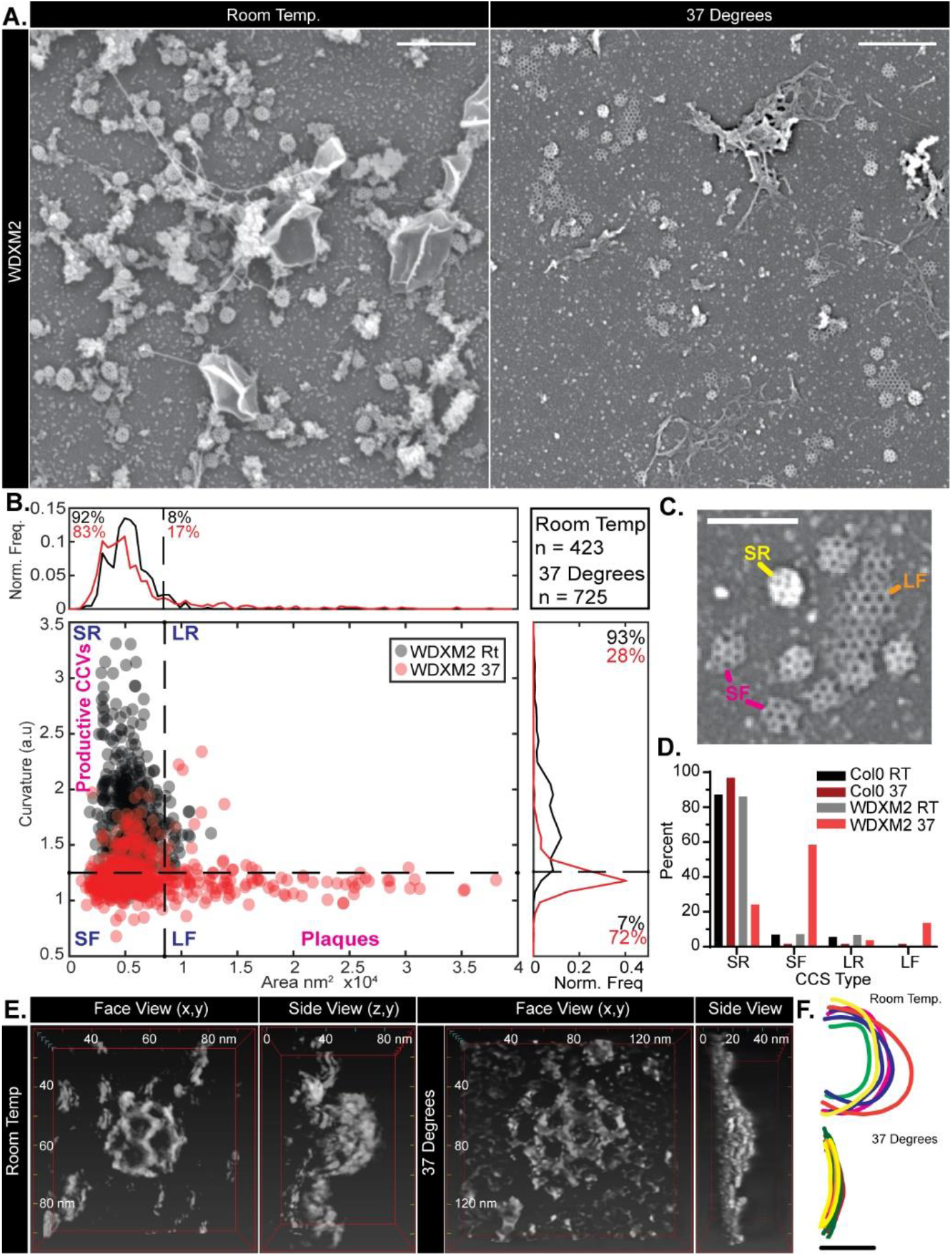
TPC disruption prevents membrane bending during plant CME. A) SEM of metal replicas of unroofed WDXM2 root protoplast cells. B) Scatter plot of the area and curvature of clathrin-coated structures (CCSs) in WDXM2 cells incubated at room-temperature (RT) (gray dots) or 37°C (red dots) for 4 hours. The graph is divided into 4 sections in order to classify the CCSs based on their shape: SR (small and round), SF (small and flat), LR (large and round) and LF (large and flat, plaques). C) Example CCSs of these classifications. D) Percentage populations of these classifications in wild type (Col-0) and WDXM2 cells subjected to RT or 37°C incubations. Data pooled from multiple experiments; N = Col-0 RT, 3 and 588 CCSs; Col-0 37°C, 3 and 127 CCSs; WDXM2 RT, 6 and 423 CCSs; WDXM2 37°C, 3 and 725 CCSs. E) Reconstructions of example STEM tomograms of clathrin structures in unroofed WDXM2 cells incubated at either RT or 37°C. F) Tracings of reconstructions overlaid each other. N = RT, 6; 37°C, 8. Scale bars, A, 500 nm; C, 200 nm; F, 50 nm

To further confirm this, we directly examined the 3-dimentional shape of clathrin structures in WDXM2 cells incubated at either control or TPC disruptive conditions using scanning transmission electron microscopy (STEM) tomography (Fig. 3E). Under conditions that disrupted TPC function, the curvature of clathrin structures did not exceed 10 nm, whereas under control conditions, the clathrin structures were spherical with Z heights > 50 nm (Fig. 3E and Movie S2-S3). Together, these ultrastructural examinations of clathrin structures demonstrate that the TPC is required to generate spherical CCVs.

Given this strong phenotype of flattened clathrin structures during TPLATE disruption, we looked for protein domains within the TPC which could mediate membrane bending. The plant-specific members of the TPC, AtEH1/Pan1 and AtEH2/Pan1, each contain 2 Eps15 homology domains (EH) which are also present in proteins which localize at the rim of CME events and known to have membrane bending activity in other systems (e.g., Eps15/Ede1 and Intersectin/Pan1) (*12, 15-17*)). As the EH domains of Eps15 have been shown to tubulate membranes *in vitro* (*17*), and the EH domains of AtEH1/Pan1 have been shown to bind membranes (*18*), we therefore tested their ability to bend membranes. To do this, we incubated large unilamellar liposomes (LUVs) with the purified EH domains of AtEH1/Pan1, and analysis by transmission electron microscopy revealed that after 2 and 30 minutes, both EH domains produced significant levels of membrane ruffling and longer tubules of vesiculated membrane compared to control treatments (Fig. 4 and Fig. S5). This demonstrated that AtEH1/Pan1 has the capacity to contribute to membrane bending. However, as AtEH1/Pan1 remains associated with the PM during WDXM2 disruption (*20*), but is not sufficient to generate the invagination (Fig. 3), it suggests that additional factors are required to modulate the membrane bending activity of the AtEH1/Pan1 EH domains. This is similar to Eps15, where the isolated EH domains can deform membranes while the full-length protein requires the co-factor FCHo (*17*). Interestingly, the Eps15 and FCHo interaction is mediated by the µ-homology domain (HD) domain within Fcho (*22*), which is similar to the interactions within the TPC, as the TML µ-HD domain interacts with AtEH1/Pan1 (*12, 18*). Although not experimentally addressed yet in the context of WDXM2, the failure to generate spherical CCVs despite AtEH/Pan1 remaining on the PM might point to the necessity of TML to also localize on the PM and likely implicates it as co-factor in plant membrane bending. Furthermore, as the µ-HD domain is absent from the TPC homologous TSET complex in *Dictyostelium*, which also lacks the AtEH/Pan1 proteins (*11, 19*), it suggests a potential functional divergence between the plant TPC and the *Dictyostelium* TSET complex, where CME is also mechanistically distinct; as in contrast to plant CME, *Dictyostelium* CME is coupled with actin (*23*).

**Fig. 4.**
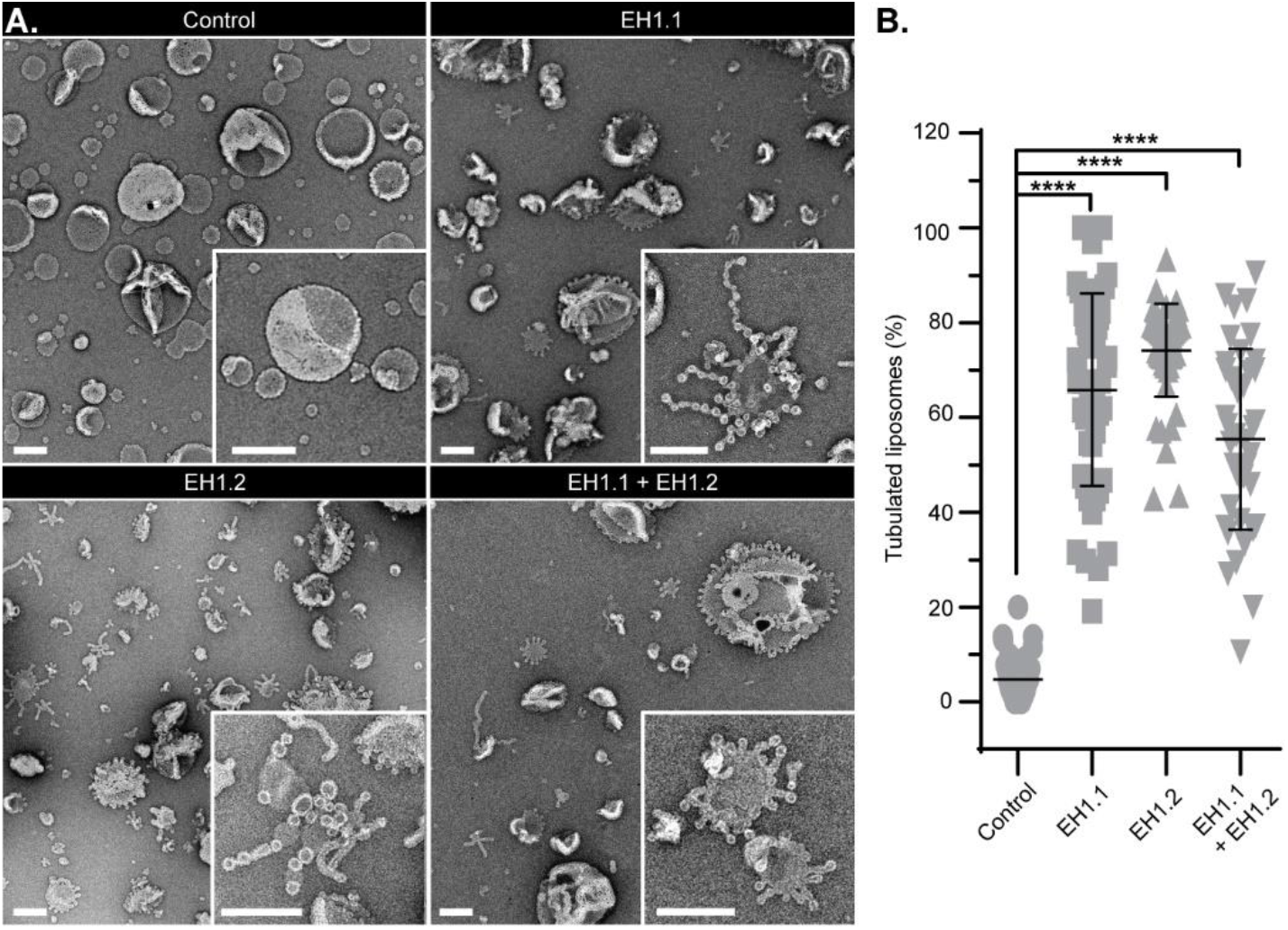
The AtEH1/Pan1 EH domains have membrane bending activity. A) Example TEM overviews of LUVs after 2 minutes incubation in control conditions, or with EH domain EH1.1, EH1.2 and EH1.1 plus EH1.2. Inserts are zooms of representative LUVs. Scale bars, A, 200 nm. B) Quantification of the percentage of LUVs which displayed tubulation. N, control, 40; EH1.1, 47; EH1.2, 52 and EH1.1+EH1.2, 40 images pooled from 3 independent experiments. Plot, mean ± SD. **** p < 0.001, One-way ANOVA with Dunnett post-test to compare to control.

**Table 1 –.**
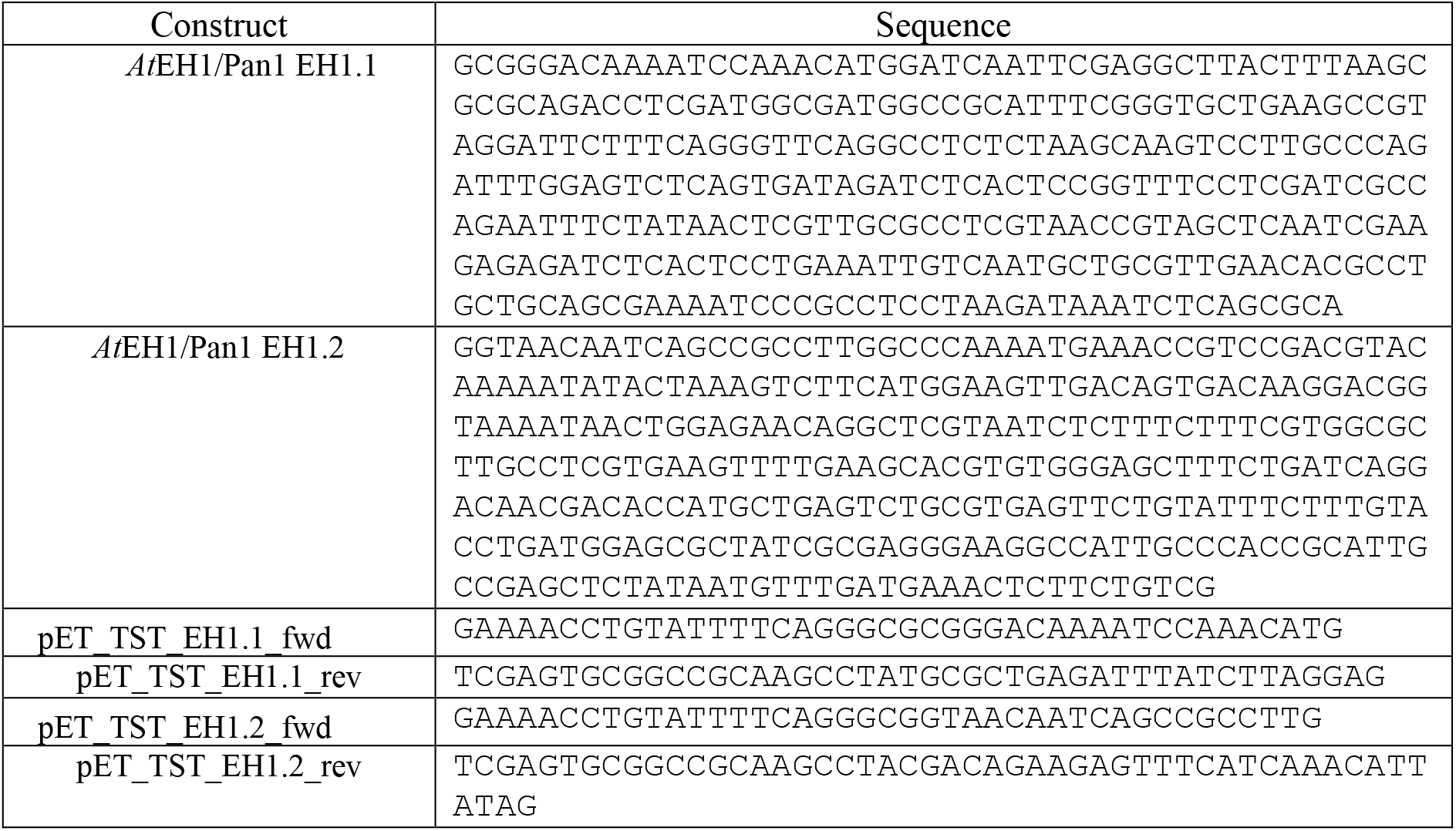
Codon optimized AtEH1/Pan1 EH domain sequences and primers used.

Overall, this study provides crucial insights into the long-standing question how endocytic vesicles can be formed by PM invaginations against the high turgor pressure of plant cells. This mystery has been used previously as a main argument against the existence of efficient endocytosis in plants and has been reinforced by the recent demonstration that, in contrast to yeast, plants do not use actin cytoskeleton to generate the necessary force for this process (*7*). Using a range of biochemistry, advanced high-, super- and ultra-resolution microscopy and *in vitro* reconstitution assays, we redefined the role of the plant-specific TPC, which has been thought to act as adaptor inside of the coat of the forming endocytic vesicles. Instead, we show that the TPC localizes outside of CCVs and is essential for membrane bending and endocytic vesicle formation (Fig. S6). This provides further evidence for the evolutionary distinct mechanism of how endocytosis operates in plants.

## Material and Methods

### Plant Materials

*Arabidopsis thaliana* accession codes for genes used in this study; AP2A1 (AT5G22770), CLC2 (AT2G40060), TPLATE (AT3G01780) and AtEH1/Pan1 (AT1G20760). Transgenic *Arabidopsis thaliana* plants used in this study were *tplate* pLAT52p::TPLATE-GFP x pRPS5A::CLC2-tagRFP, pLAT52p::TPLATE-GFP x pRPS5A::AP2A1-TagRFP (*8*) and *tplate* pLAT52::WDXM2-GFP (*20*).

### Growth Conditions

Plants are grown by plating seeds on to 1/2 AM agar plates with 1% (w/v) sucrose, stratified for 2-3 days in the dark at 4°C, then transferred to growth rooms (21°C, 16 hours light, 8 hours dark) and grown vertically for 4, 5 or 7 days as detailed below.

### Departure Analysis of EAPs from CCVs

Raw data from Wang et al. (*9*) was analyzed to determine the departure dynamics of the endocytosis proteins. Briefly, spinning disk microscopy was conducted on 4-day old epidermal cells of etiolated hypocotyls were imaged with a Nikon Ti microscope equipped with a Ultraview spinning-disk system (Perkin-Elmer), a Plan Apo 100x 1.45 NA oil immersion objective and a CherryTemp system (Cherry Biotech) to apply the experimental temperature conditions; either room temperature (25°C) or 12 °C. Time lapses were collected at a frame rate of 1 frame per 1.174s. The 12°C time-lapses of TPLATE-GFP and CLC2-TagRFP samples were subjected to histogram matching bleach correction and then dynamically re-sliced to produce kymographs in Fiji (*24*). The CLC2 channel was manually screened to identify kymograph traces with a visible departure track. These selected traces were then examined to compare the departure of both channels and categized as illustrated in Fig. S1.

### Western blotting analysis of CCV purification

CCVs were purified from suspension cultured Arabidopsis T87W cells as previously described (*25*). Equal amounts of protein from the deuterium ficoll gradient load (DFGL) and purified clathrin coated vesicle (CCV) samples were separated by SDS-PAGE, transferred to nitrocellulose membrane, and immunoblotted with anti-CLC2 1:10,000 (*26*), anti-CHC 1:1,000 (sc-57684, Santa Cruz Biotechnology), anti-AP2mu2 1:250 (*27*), anti-TPLATE 1:2000 (*28*), and anti-DRP1c 1:500 (*29*) antibodies. Primary antibodies were detected through anti-rabbit or anti-mouse secondary antibodies (Sigma Aldrich) conjugated to horseradish peroxidase at 1:5,000 before application of SuperSignal West Femto enhanced chemiluminescent (ECL) substrate (ThermoFisher) and subsequent imaging with iBright CL1000 Imaging System (ThermoFisher). The integrated density values of the chemiluminescent bands of the DFGL and CCV fractions were measured by ImageJ (NIH). The integrated density value of the CCV band was divided by the corresponding value of the DFGL band to determine the relative enrichment across three independent CCV purifications.

### FM uptake and TIRF-M imaging and analysis

A Zeiss LSM-800 confocal microscope was to examine the effect of FM4-64 uptake in 5-day old Col-0 seedlings. Seedlings were incubated for 6 hours at either room temperature or 35°C for six hours and then incubated with 2 µM FM4-64 in AM+ media for 5 minutes, washed twice in 1/2 MS and 1% sucrose media and imaged and analyzed as specified previously (*30*). A 40x water immersion objected was used.

Total internal reflective fluorescence microscopy (TIRF-M) experiments made use of an Olympus IX83 inverted microscope equipped with a Cell^TIRF module using an OLYMPUS Uapo N 100x/1.49 Oil TIRF objective. 7-day old seedlings were incubated at either 25°C or 37°C for 6 hours prior to imaging. Root epidermal cells were imaged and analyzed as described previously (*30*); this provided unbiased lifetimes, densities and fluorescence profiles of endocytosis proteins in samples subjected to the experimental temperature conditions.

### Super-resolution localization of EAPs

Structured illumination microscopy (SIM) was conducted on 7-day old seedlings expressing TPLATE-GFP or AP2A1-GFP and CLC2-TagRFP. Root samples were prepared as described previously (*30*), but high-precision 1.5 coverslips were used (Thorlabs, #CG15CH), and epidermal cells in the elongation zone were selected for imaging. For 3D-SIM, an OMX BLAZE v4 SIM (Applied Precision) was used. For TIRF-SIM, an OMX SR (GE Healthcare) was used. Both are equipped with a 60 × 1.42 NA oil immersion objective, and 100 mw 488 nm and 561 nm lasers were used for illumination (for TPLATE-GFP x CLC2-TagRFP, 488 laser powers ranged from; 488, 30-100%; 561, 25-100%. For TPLATE-GFP x AP2A1-TagRFP, laser powers ranged from; 488, 20-100%; 561, 40-100%). Images were reconstructed using SOFTWORX (GE Healthcare) and further processed in Fiji (*24*).

The co-localization rate was determined by using ComDet (https://github.com/ekatrukha/ComDet), where co-localization was determined positive if spot detection were less than 4 pixels apart. This method uses wavelet decomposition to determine spot detection, thus considers rings and spots extremely close together as a single spot. To determine the pattern of localization of TPLATE, i.e., if it is a spot or surrounding the CME event, spots were manually examined and scored if TPLATE presented as a crescent or ring around a CLC2 or AP2A1 spot.

### Ultra-structural examination of CCVs by SEM and EM tomography from metal replicas of protoplasts made directly from roots

Densely sown Col-0 or WDXM2-GFP plants were grown for 8-10 days. The roots were cut into small ∼1-2 mm fragment directly in to ‘Enzyme solution’ (0.4 M Mannitol, 20 mM KCl, 20 mM MES pH 5.7, 1.5% Cellulase R10 (Yakult), 0.4% Macerozyme R10 (Yakult) in H20). The cuttings and enzyme solution were placed into a vacuum chamber for 20 mins and then subjected to a 3-hour incubation at room temperature, in the dark, and with gentle agitation. The cells were then centrifuged at 100 rcf for 2 mins, and the pellet washed with ‘W5 buffer’ (154 mM NaCl, 125 mM CaCl2, 5 mM KCL, 2mM MES) by centrifugation (100 rcf for 2 mins). The cells were then resuspended in W5 buffer and incubated at 4°C for 30 mins. The sample was again centrifuged at 100 rcf for 2 mins, and the cells were resuspended in ‘hyperosmotic GM buffer’ (GM; 0.44% (w/v) Murashige and Skoog (MS) powder with vitamins (Duchefa Biochemie), 89 mM Sucrose, 75 mM mannitol, pH 5.5 adjusted with KOH) and then plated on precleaned (washed in pure ethanol and sonicated), carbon (10 nm thickness) and poly-l-lysine (Sigma) coated coverslips. Samples were incubated at room temperature in the dark for 30 mins, and then subjected to a 4-hour incubation in the dark at either room temperature or 37°C. Samples were then unroofed as described previously (*30*), with the buffers were equilibrated to either room temperature or 37°C. Samples for SEM analysis were attached to SEM mounts using sticky carbon tape and coated with platinum to a thickness of 3 nm, whereas samples for STEM were attached to a sticky Post-It-Note (as described in (*31*)) and coated with 3 nm platinum and 4 nm carbon using an ACE600 coating device (Leica Microsystems). The STEM samples were then washed with Buffered Oxide Etchant (diluted 6:1 with surfactant) to separate the metal replica from the coverslip, washed with distilled water, and remounted on formvar/carbon-coated 200-line bar EM grids (Science Services).

The SEM samples were then imaged with an FE-SEM Merlin Compact VP (Zeiss) and imaged with an In-lens Duo detector (SE and BSE imaging) at an accelerating voltage of 3-5 kV. The area and mean grey value of clathrin-coated structures (CCSs) was measured using Fiji (*24*), where ROIs were manually draw around each CCS. To estimate the curvature of the CCSs, the mean CCS ROI was divided by the average grey value of the PM (as determined by the mean grey value of 4 PM ROIs in each corner of each image). From these two values, the morphology of the CCS could be determined by using thresholds and divided into categories, as described by *Moulay et al*., (*21*). We used an area threshold of 8500 mm^2^ (which is derived from a diameter of 105 nm) to determine if the CCS was small or large, and a curvature value of 1.25 (determined by measuring the mean grey value of the large CCSs observed in TPC disruption conditions) to determine if the CCS was round or flat. Pooled data from multiple experiments were plotted and the percentage of CCVs in each category was calculated.

STEM Tomograms were recorded using a JEOL JEM2800 scanning/transmission electron microscope (200kV). Each CCV was imaged at over a range of -72° to 72°, with 4° steps driven by STEM Meister (TEMography.com). Tomograms were then processed and 3D reconstructions were made using Composer and Evo-viewer (TEMography.com). To examine the curvature, 3D reconstructions were rotated 90° and their profiles were manually traced in Adobe Illustrator.

### Expression and purification of AtEH/Pan1 EH domains

The two EH domains of atEH1, EH1.1 and EH1.2 as defined by (*18*), were amplified from synthetic AtEH1/Pan1 (codon optimized for bacterial expression, IDT) (Supplemental Table 1) and inserted into pET-TwinStrep-TEV-G4. They were then expressed in *E. coli* BL21 cells and grown at 37°C in Lysogeny Broth medium (pH 7.0) supplemented with 50 µg ml-1 kanamycin. Protein expression was induced at an OD600 of 0.6 with 1 mM isopropyl-β-thiogalactopyranoside (IPTG) and incubated for 5 h at 37°C. Cultures were centrifuged at 5000 g for 30 min at 4°C, and pellets were resuspended in 50 mL Phosphate-buffered saline (PBS) buffer. They were then centrifuged at 4700 g for 30 min at 4°C and then pellets were frozen and stored at –80°C until further processing.

The pellets were resuspended for 1 hour at 4°C with gentle mixing in buffer A (20 mM HEPES (pH7.4), 150 mM NaCl, 2 mM CaCl2, as described by (*18*)) with supplemented ethylenediaminetetraacetic acid (EDTA)-free protease inhibitor cocktail tablets (Roche Diagnostics), 1 mM phenylmethylsulfonyl fluoride (PMSF), 1 mg mL-1 lysozyme, 1 µg ml-1 DNase I. Cells were lysed by sonication (Qsonica Q700) and centrifuged at 67000 g for 1 h at 4 °C. The clarified lysate was incubated with Streptactin sepharose resin (Strep-Tactin® Sepharose® resin; iba) for 1 hour at 4°C. The resin was washed with 40 bed volumes of buffer A and the fusion protein was eluted with buffer A containing 5 mM d-Desthiobiotin (Sigma-Algrich). Peak atEH domain fractions were dialyzed overnight at 4°C against buffer A in the presence of TwinStrep-tagged TEV protease (*32*) at a protease-to-sample molar ratio of 1:100. After centrifugation (21 140 xg for 10 min at 4 °C), the supernatant was applied to a HiLoad 16/600 Superdex 75 pg column, pre-equilibrated with buffer A, using a fast protein liquid chromatography system. Protein was eluted with buffer A and stored in aliquots at –80 °C. The protein sequences of the EH domains were verified by MS analysis.

### Liposome tubulation assay

LUVs were prepared using a mixture of 1,2-dioleoyl-sn-glycero-3-phospho-(1’-rac-glycerol) (DOPC), 1,2-dioleoyl-sn-glycero-3-phospho-L-serine (DOPS), cholesterol (plant) and 1,2-dioleoyl-sn-glycero-3-phospho-(1’-myo-inositol-4’,5’-bisphosphate) (PI(4,5)P2) (Avanti) at a ratio of 60:17.5:20:2,5 mol% was used. Lipids were mixed in a glass vial at the desired ratio, blow-dried with filtered N2 to form a thin homogeneous film and kept under vacuum for 2–3 h. The lipid film was rehydrated in a swelling buffer (20 mM HEPES (pH 7.4), 150 mM NaCl) for 10 min at room temperature to a total lipid concentration of 2 mM. The mixture was vortexed rigorously and the resulting dispersion of multilamellar vesicles was repeatedly freeze–thawed (5–6 times) in liquid N2. The mixture was extruded through a polycarbonate membrane with pore size 400 nm (LiposoFast Liposome Factory). LUVs were stored at 4 °C and used within 4 days. To assay the membrane bending activity of proteins of interest upon the LUVs, 10 µM of the protein of interest was mixed with 0.5 mM of LUVs in swelling buffer and incubated for 2, or 30, minutes at room temperature. Control LUVs were diluted to a concentration of 0.5 mM in swelling buffer and incubated for 1 hour at room temperature. 20 ul of the experimental solutions were incubated on glow-discharged to carbon-coated copper EM grids (300 mesh, EMS). Filter paper was used to remove any excess solution and the EM samples were then washed three times with swelling buffer. They were then negatively stained with 2% uranyl acetate aqueous solution for 2 mins and observed under a Tecnai 12 transmission electron microscope operated at 120 kV (Thermo Fisher Scientific). The number of tubulated and non-tubulated liposomes was counted manually using Fiji (*24*) from multiple experiments.

## Acknowledgments

We gratefully thank Julie Neveu and Dr. Amanda Barranco of the Grégory Vert lab for helping preparing plants in France. Dr. Zuzana Gelova for help and advice with protoplast generation. Dr. Stéphane Vassilopoulos and Dr. Florian Schur for advice regarding EM tomography. Alejandro Marquiegui Alvaro for help with material generation. This research was supported by the Scientific Service Units (SSU) of IST-Austria through resources provided by the Electron microscopy Facility (EMF), Lab Support Facility (LSF) (particularly Dorota Jaworska) and the Bioimaging Facility (BIF). We acknowledge the Advanced Microscopy Facility of the Vienna BioCenter Core Facilities for use of the 3D SIM. For the Mass Spectrometry analysis of protein, we acknowledge BOKU Core Facility Mass Spectrometry. And Lukasz Kowalski for generously gifting us mWasabi.

## Author contributions

Conceptualization: A.J. and J.F.; Investigation: A.J., D.A.D, N.G., W.A.K., V.Z., T.C., P.M., M.H., and J.S.; J.W. material generation. Formal Analysis: A.J., D.A.D and N.G.; Writing – original draft: A.J. and J.F.; Writing – review & editing: all authors; Supervision: D.V.D., E.B., M.L., S.Y.B. and J.F.

## Supplemental Figures

**Fig. S1.**
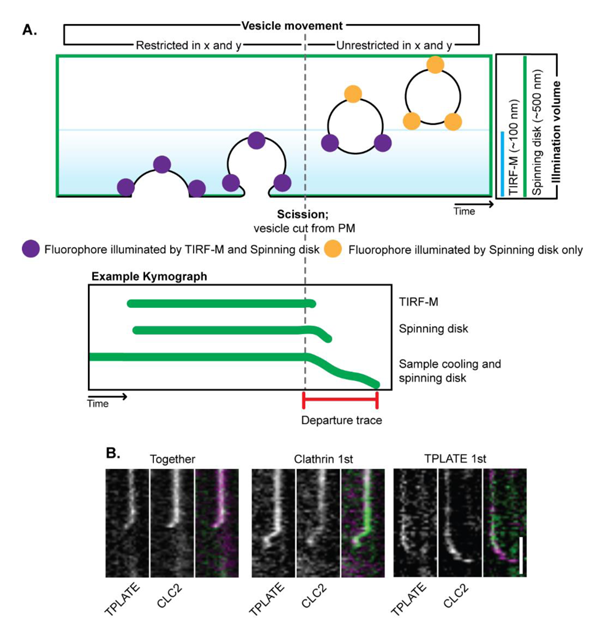
Classification of departure events. Related to figure 1. A) During CME, as the CCV invaginates on the PM it is restricted in movement in X and Y. This means in a kymograph it will present a linear pattern. However, once it is freed from the PM (gray dotted line), it trafficks away from the PM and thus is able to move in all directions, resulting in a lateral movement in the kymograph trace. Notably, this also results in the CCV moving deeper into the cell and exiting the illumination volume of fluorescence imaging methods, which determine how long the CCV is visible once after being cut from the PM and thus how long the departure trace is on a kymograph. TIRF-M has a Z depth of around 100-200 nm, whereas spinning disc has a Z volume of ∼500 nm in Z away from the PM, which means fluorescently labeled CCVs are visible longer with spinning disk and thus have a longer departure trace. This is extended by using sample cooling to slow down cellular dynamics. B) Example departure traces of TPLATE (green) and CLC2 (magenta) at single events of CME demonstrating 4 different types of departure. ‘Together’, where both traces display the same lateral movement at the end of the trace and disappear together, which would be expected if TPLATE is bound to the CCV under CLC2 as it cannot departure the CCV before uncoating occurs before leaving the illumination volume; ‘Clathrin 1^st’^, where the CLC2 trace disappears before the TPLATE trace, which could indicate that CLC is removed as a layer before TPLATE and ‘TPLATE 1^st’^, where the TPLATE trace terminates before the CLC2 trace, indicating that it is free to leave the CCV before the clathrin coat dissociates. Scale bar, B, 60 s.

**Fig. S2.**
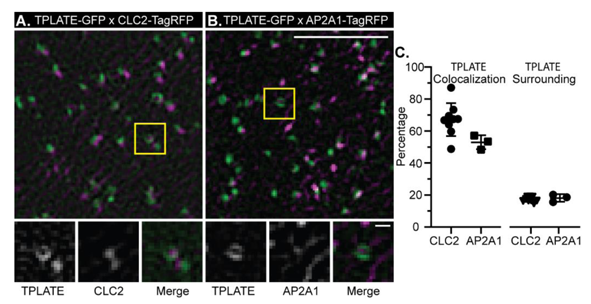
TPLATE localizes around Clathrin and AP2. Related to figure 2. TIRF-SIM example images of *Arabidopsis* root epidermal cells expressing A) TPLATE-GFP (green) x CLC2-TagRFP (magenta) and B) TPLATE-GFP x AP2A1-TagRFP. Yellow squares denote the area zoomed in for the lower panels. C) Quantification of colocalization of TPLATE with CLC2 and AP2A1, and the percentage of colocalized spots where TPLATE is found to surround CLC2 and AP2A1. Plots, mean ± SD. N; TPLATE x CLC2, 9 cells; TPLATE x AP2, 3 cells. Scale bars, A and B, upper image, 2 µm, lower panels, 200 nm.

**Fig. S3.**
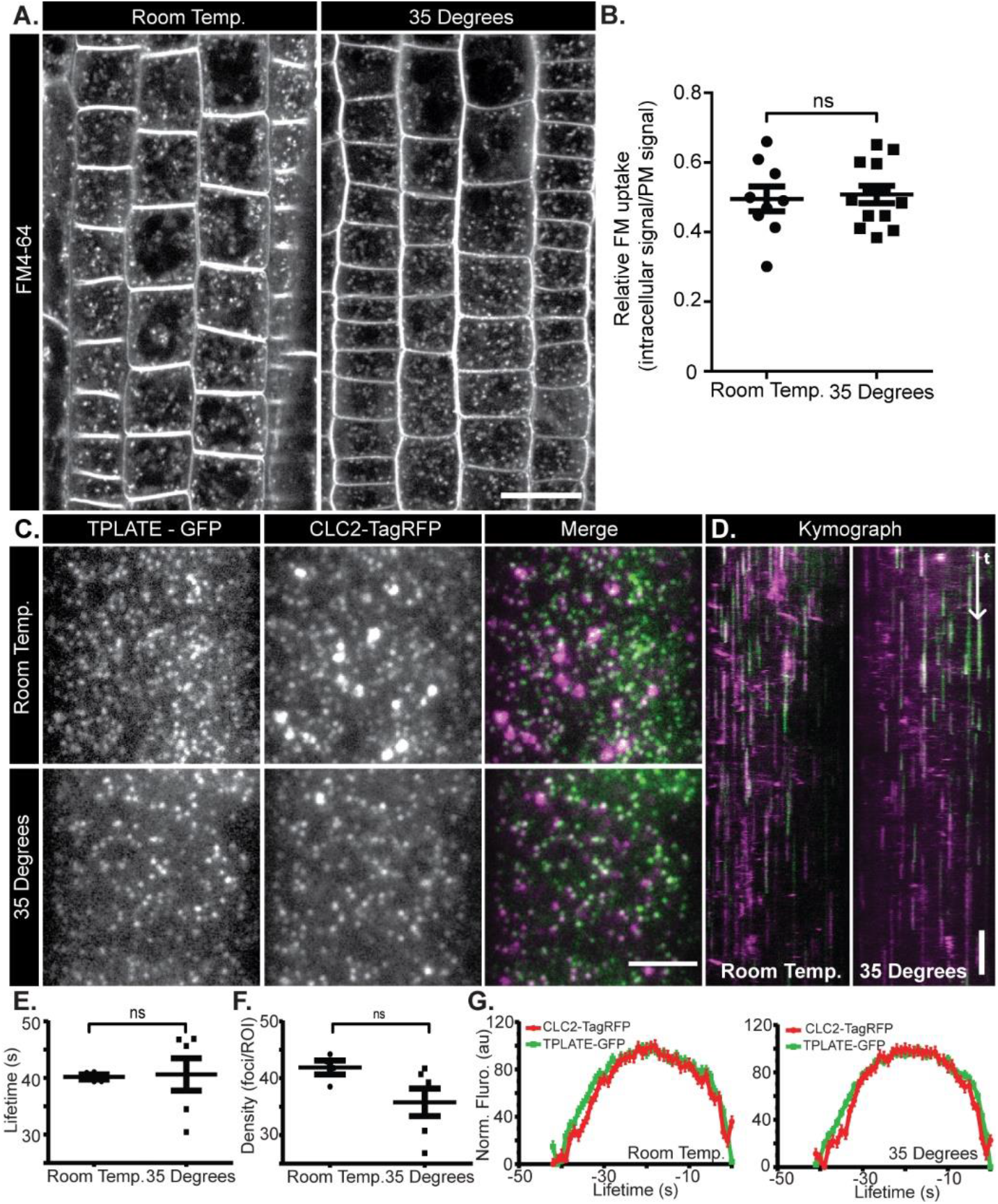
Heat shock does not affect the efficiency and dynamics of endocytosis. Related to figure 3. A) Confocal images of Arabidopsis roots incubated with FM4-64 after a 6-hour incubation at room temperature or 35°C. B) Quantification of FM uptake from multiple experiments. N; room temperature, 9 independent seedlings, 169 cells; 35°C, 13 independent roots, 245 cells. C) Example TIRF-M images of Arabidopsis root epidermal cells expressing TPLATE-GFP (green) and CLC2-TagRFP (magenta) after incubation at room temperature or 35°C for 6 hours. D) Typical kymographs obtained from data shown in C. E) The lifetimes, F) density and G) mean profiles of endocytosis events combined for multiple experiments of seedlings incubated at room temperature or 35°C for 6 hours. N; room temperature, 4 independent roots, 18473 events; 35°C, 6 independent roots, 22223 events. Ns = not significant (P > 0.05) t-test result. Scale bars, A, 20 µm; C, 5 µm; D, 60 s.

**Fig. S4.**
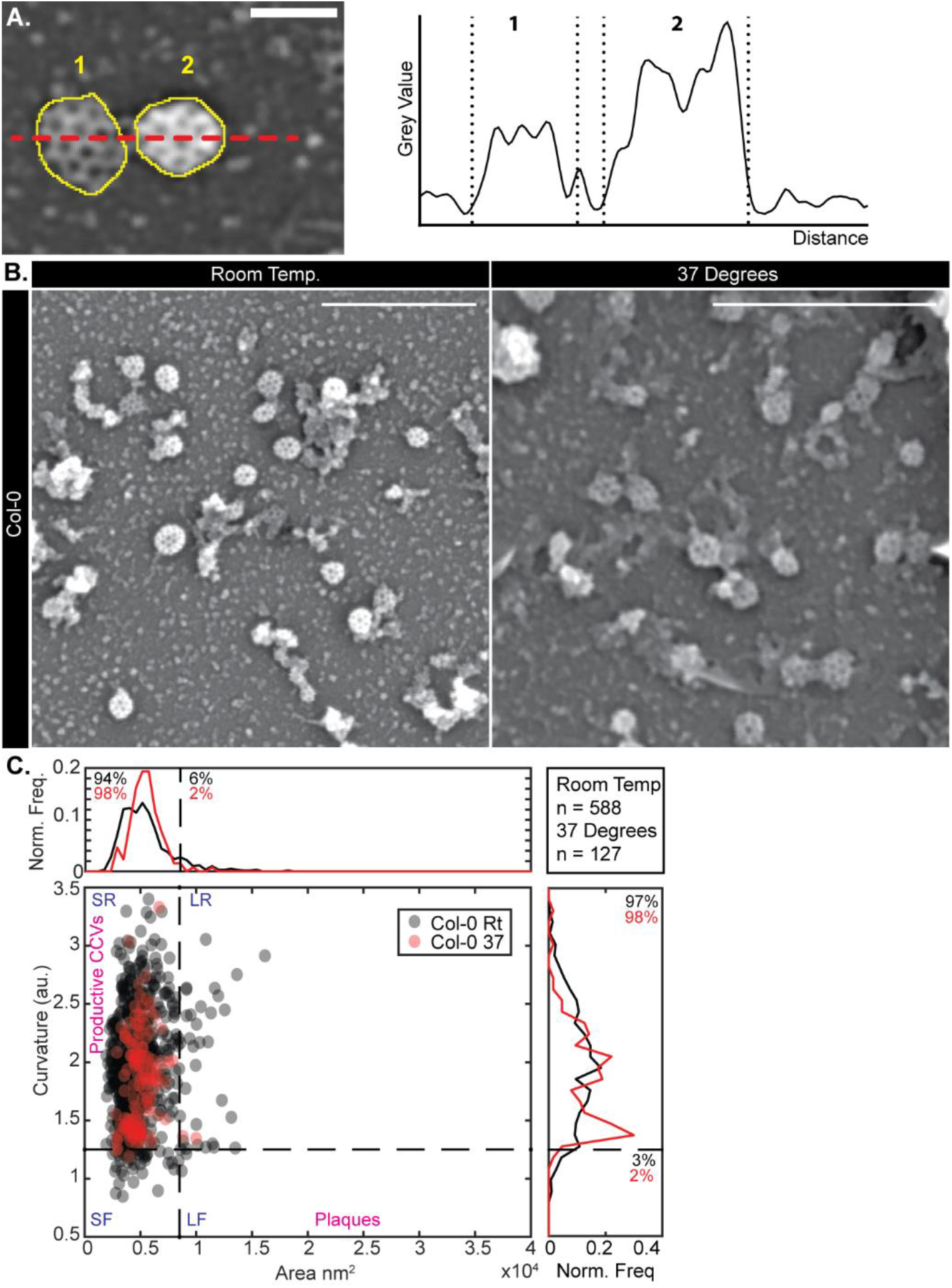
Heat shock does not affect the formation of spherical clathrin-coated vesicles. Related to Figure 3. A) ROIs (yellow dashed line) are drawn around the clathrin-coated vesicles (CCS) to determine the size and the mean gray values. The gray values can be used to estimate the curvature of the CCSs, for example, the flatter CCS (1) has a lower grey value than the rounder CCS (2). B) Representative SEM images of metal replicas of unroofed Col-0 root protoplast cells. C) Scatter plot of the area and curvature of CCSs in wild-type Col-0 cells incubated at room temperature (RT) (gray dots) or 37°C (red dots) for 4 hours and classified into 4 types of CCSs as detailed in the methods. Scale bars, 500 nm. N = RT, 3 and 588 CCSs; 37°C, 3 and 127 CCSs. Scale bars, A, 100 nm; B, 500 nm.

**Fig. S5.**
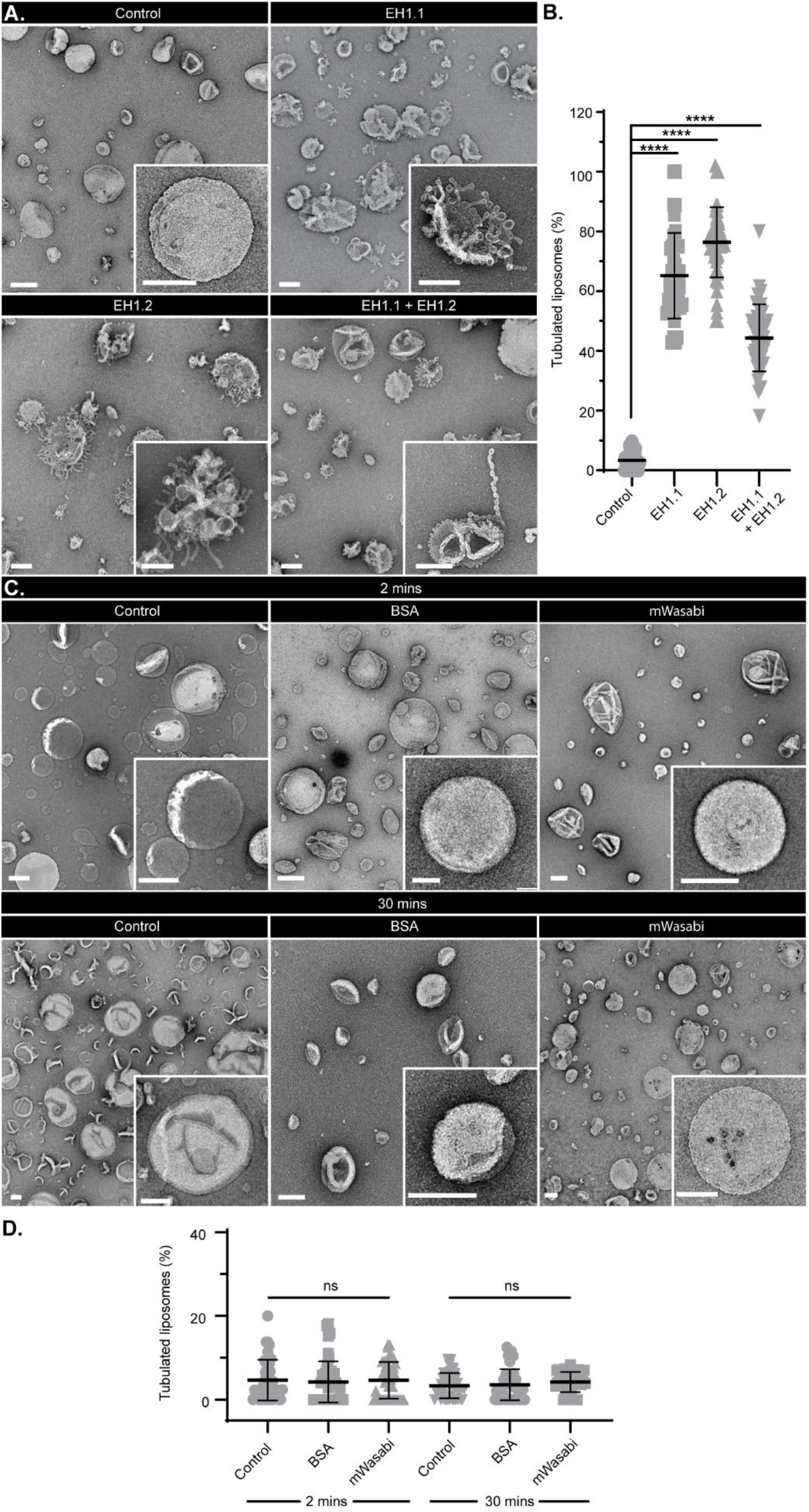
AtEH1/Pan1 EH domain membrane bending activity and additional controls. Related to figure 4. A) Example TEM overviews of LUVs after 30 minutes incubation in control conditions, or with EH domain EH1.1, EHD1.2 and EHD1.1 plus EHD1.2. Inserts are zooms of representative LUVs. B) Quantification of the percentage of LUVs which displayed tubulation. N; control, 41; EH1.1, 40; EH1.2, 49 and EH1.1+EH1.2, 46 images pooled from 3 independent experiments. C) Example TEM overviews of LUVs after 2 minutes (upper panels), or 30 mins (lower panels) incubation in control conditions, or with BSA, or mWasabi. Inserts are zooms of representative LUVs. D) Quantification of the percentage of LUVs which displayed tubulation. N; control 2 min, 41; BSA 2 min, 48; mWasabi 2 min, 42; control 30 min, 4; BSA 30 min, 42 and mWasabi 30 min, 40 images pooled from 3 independent experiments. Plot, mean ±SD. **** p < 0.001, one-way ANOVA with Dunnett post-test to compare to control, ‘ns’ indicate not significant. Scale bars, 200 nm.

**Fig. S6.**
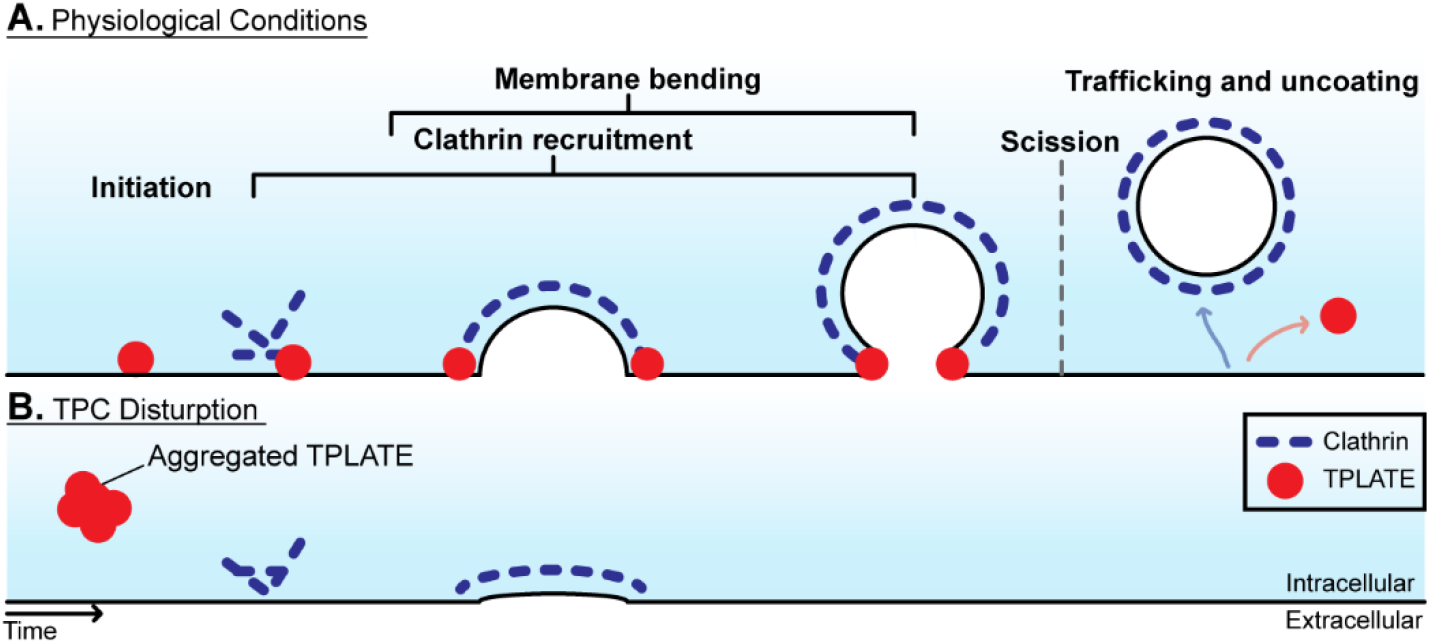
The TPC is required for membrane bending during plant CME. A) In physiological conditions TPLATE and the TPC is recruited to the PM and clathrin is then recruited to begin the coat assembly phase of CME. Then the membrane bends creating an invagination against the high turgor pressure, here TPLATE is located at the rim of this invagination and drives the membrane bending. Eventually as the CME vesicle grows, a tight neck is created, where the TPC ring closes and after scission is able to departure the CCV before clathrin as it is not associated within the coat. B) During TPC disruption, TPLATE aggerates in the cytoplasm, while clathrin and AtEH1/Pan1 is still recruited to the PM but no invagination is created as there the full TPC is not present, resulting in the failure of membrane bending during the CME event.

**Movie S1. TIRF-SIM of root epidermal cells expressing TPLATE-GFP**

Example TIRF-SIM time lapse of TPLATE-GFP in an *Arabidopsis* root epidermal cell. Yellow ovals denote TPLATE structures which dynamically form ring structures. Time interval between frames, 5 seconds. Scale bar, 5 µm.

**Movie S2. STEM tomography of a clathrin-coated vesicle in control conditions**

180° rotation of a 3D reconstructed STEM tomogram for a CCV in a metal replica of an unroofed WDXM2 cell after 4-hour incubation at room temperature.

**Movie S3. STEM tomography of a clathrin-coated vesicle during TPC disruption**

180° rotation of a 3D reconstructed STEM tomogram for a CCV in a metal replica of an unroofed WDXM2 cell after 4-hour incubation at 37°C.

